# Genetic Variation in Organ System Decline Across the *Drosophila* Genetic Reference Panel

**DOI:** 10.64898/2026.09.14.750524

**Authors:** James F. Del Tito, Lesley A. Alton, Matthew D. W. Piper, Christen K. Mirth

**Affiliations:** School of Biological Sciences, Monash University, Melbourne, Victoria, 3800, Australia

## Abstract

Genetic background has a significant impact on an organism’s lifespan. Underlying this metric, ageing is characterised by a sequence of physiological systems decline that leads to death. Despite this, much of the ageing research performed on model organisms in the lab uses lifespan, rather than physiological decline, as a metric to measure ageing. Here, we sought to establish if differing genotype alters age-related organ system decline by repeatedly measuring the function of five organ systems throughout the lifespan of 19 lines from the *Drosophila* Genetic Reference Panel. We found that the magnitude of each organ system’s decline differs across genotypes, and that different organ systems decline at different rates within a genotype. We also uncovered that while declines in activity rate and reproduction were associated with length of life, overall patterns of phenotypic organ system decline differed significantly across genotypes. Overall, these data suggest that individuals with different genetic backgrounds have distinct patterns of physiological decline, implying that the mechanisms driving ageing vary between individuals. These data underscore the importance of studying organ systems decline in addition to lifespan to gain a full understanding of ageing.

## Introduction

Ageing, the gradual decline in physiological function over time, is an inevitable aspect of life (1). Evolutionary theory maintains that organisms age because the force of natural selection weakens with age such that its negative effects manifest after an organism would typically die from external hazards, making it beyond the reach of natural selection (2). Ageing can therefore evolve, either through accumulation of late-acting deleterious mutations, or through links to other traits that are themselves favoured by selection, such as growth and reproduction (3–6). For this reason, ageing and its physiological decline is thought to be indirectly determined and likely to progress in a non-stereotyped fashion, differing between individuals as a function of their particular environment and genetic makeup (7).

Many ageing studies, particularly those which utilise model organisms like *D. melanogaster,* focus on population level features of ageing such as mean, median, and maximum lifespan (8). These traits are informative and are relatively easy to measure on a large scale, however they do not fully represent ageing as they do not account for the physiological decline underlying these metrics (9). Ageing is a complex process, and the decline of cells, tissues, and organ systems may occur at different rates in different individuals in ways that may not correlate with organismal lifespan (10, 11). To understand how treatments can sustain healthy ageing, it is important to better characterise how the physiological progression of ageing occurs.

Age-related organ system decline, and thus how variation in healthspan relates to lifespan, is an important area of interest within biogerontology. In particular, the role of the gut has become a focus, as age-related increases in gut permeability can be predictive of death in model organisms like the fruit fly *Drosophila melanogaster,* the nematode worm *Caenorhabditis elegans* and in zebrafish, *Danio rerio* (12–14). It has also been established that the function of other organ systems, like the reproductive and neuromuscular systems, decline with age, and that their relative rates of decline vary with genotype (15, 16). While recent studies have begun to track and compare physiological decline across multiple organ systems simultaneously, this work remains limited when compared with that measuring lifespan (17–19). Moreover, how genetic background shapes variation in patterns of multi-system decline remains largely undetermined in lab models. Identifying inter-individual differences in decline would clarify how genetic variation shapes ageing and identify a broader range of targets for research into its underlying mechanisms.

Isogenic fly lines provide a means of detecting genetic variation in a trait. The *Drosophila* Genetic Reference Panel (DGRP) are a panel of fully sequenced, inbred strains of *Drosophila melanogaster* which contain a representative sample of the segregating genetic variation found in natural populations (20). The panel has been used in many studies to identify the effects of genetic variation in age-related traits like lifespan, locomotor activity, and heat knockdown time, and as such are a useful tool to identify if there is genetic variation in overall patterns of organ decline (21–23). However, studies are yet to assess these metrics in combination to determine their relative rates of decline across genotypes.

Here, we explore how varying genetic background affects the age-related decline of various aspects of organismal function that represent organ systems. These include indicators of general metabolism, gut integrity, fat body/oenocytes function in detoxification, reproductive system decline, and the neuromuscular system’s ability to support movement in adult female *D. melanogaster* across 19 isogenic lines from the DGRP. By characterising the decline of all these indicators and how they associate with lifespan across these lines, we report the extent to which genotype modifies the progression of ageing-related system dysfunction that leads to death.

## Results

### Isogenic fly lines from the DGRP differ significantly in their lifespan

Lifespan is commonly used as a proxy measure for ageing in many studies (24, 25). To assess variation in ageing using lifespan, the trait was measured in 19 lines from the Drosophila Genetic Reference Panel (DGRP). Lines selected for this study were all from the Core 40 set (except for RAL776), which are a subset of the DGRP which represent the most genetically distinct lines found in the panel, and with sufficient viability to obtain enough flies to run the experiments (26). We found that there was a large and significant variation in lifespan across lines (Figure 1; p < 0.01). The shortest-lived line (RAL360) lived for 29.4 days on average while the longest-lived line, RAL774, lived almost twice as long with average lifespan of 52.5 days.

**Figure 1.**
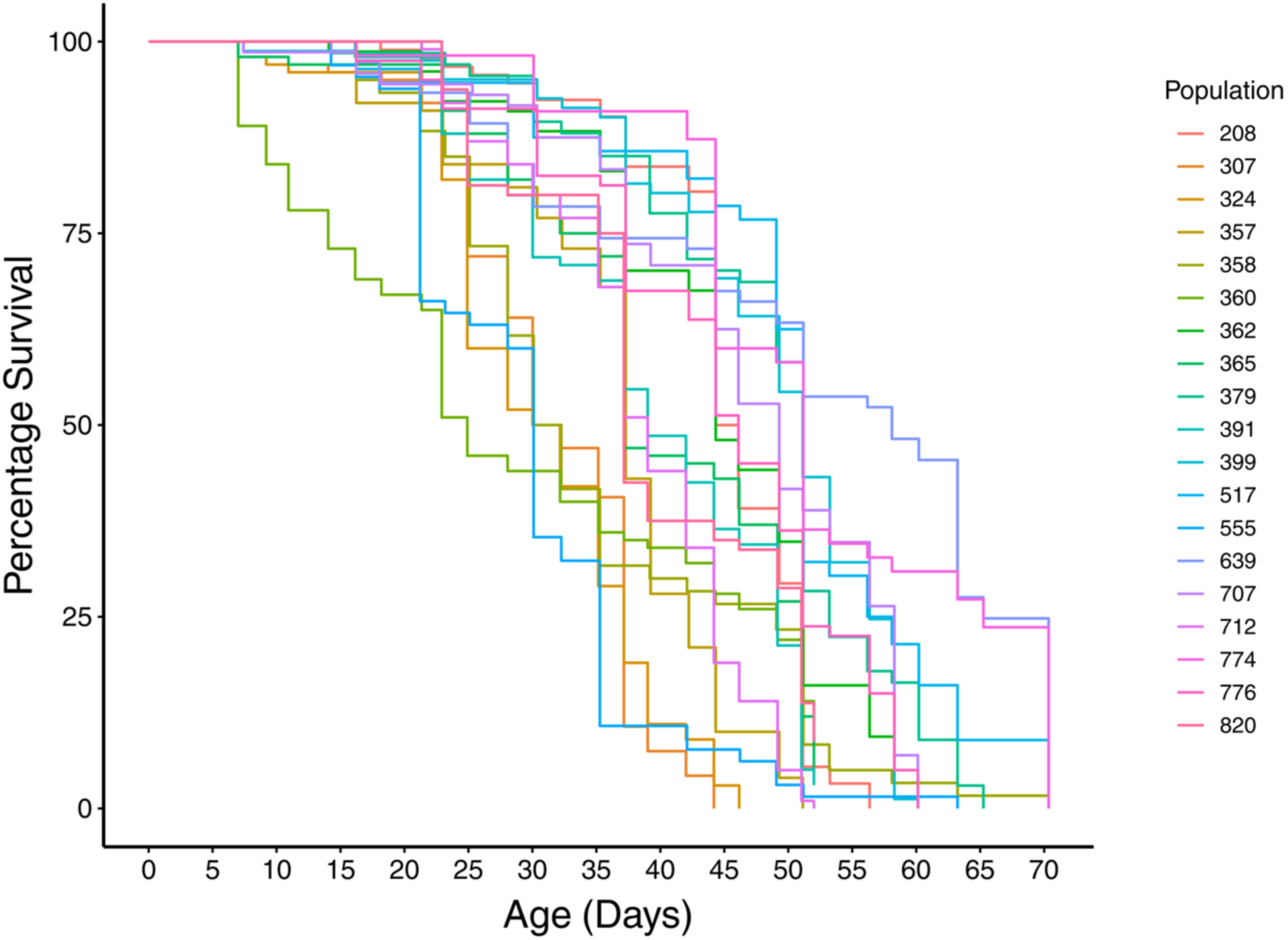
Isogenic fly lines from the DGRP differ significantly in their lifespan. Isogenic DGRP lines were maintained on a nutritionally complete (sugar / yeast) diet throughout life with deaths and censors recorded every two to three days. Lifespan differed significantly between isogenic lines (Cox Proportional Hazards: p < 0.01. n = 100 flies per line, Supplementary table S1).

### Changes in physiological parameters with age varies significantly between genotypes

While our lifespan data indicated that there is variation in ageing across DGRP lines, we wanted to assess whether this variation was also reflected in their physiological decline with age. To do this, we selected five phenotypic assays, each reflecting the function of a distinct organ system, and measured them repeatedly across the lifespan of flies from the same 19 DGRP lines used for lifespans (Figure 2a-e). The assays were egg counts, to measure reproductive output; nicotine survival assays, to measure fat body / oenocyte detoxification function (27, 28); smurf assays, to measure gut integrity (29); respirometry, to measure general metabolic function (30); and activity assays using *Drosophila* Activity Monitors (DAMs), to measure neuromuscular function (31). The averaged performance across all genotypes declined with age (p = < 0.05), except for CO_2_ production (Figure 2d, p = 0.85), which remained stable with age.

**Figure 2.**
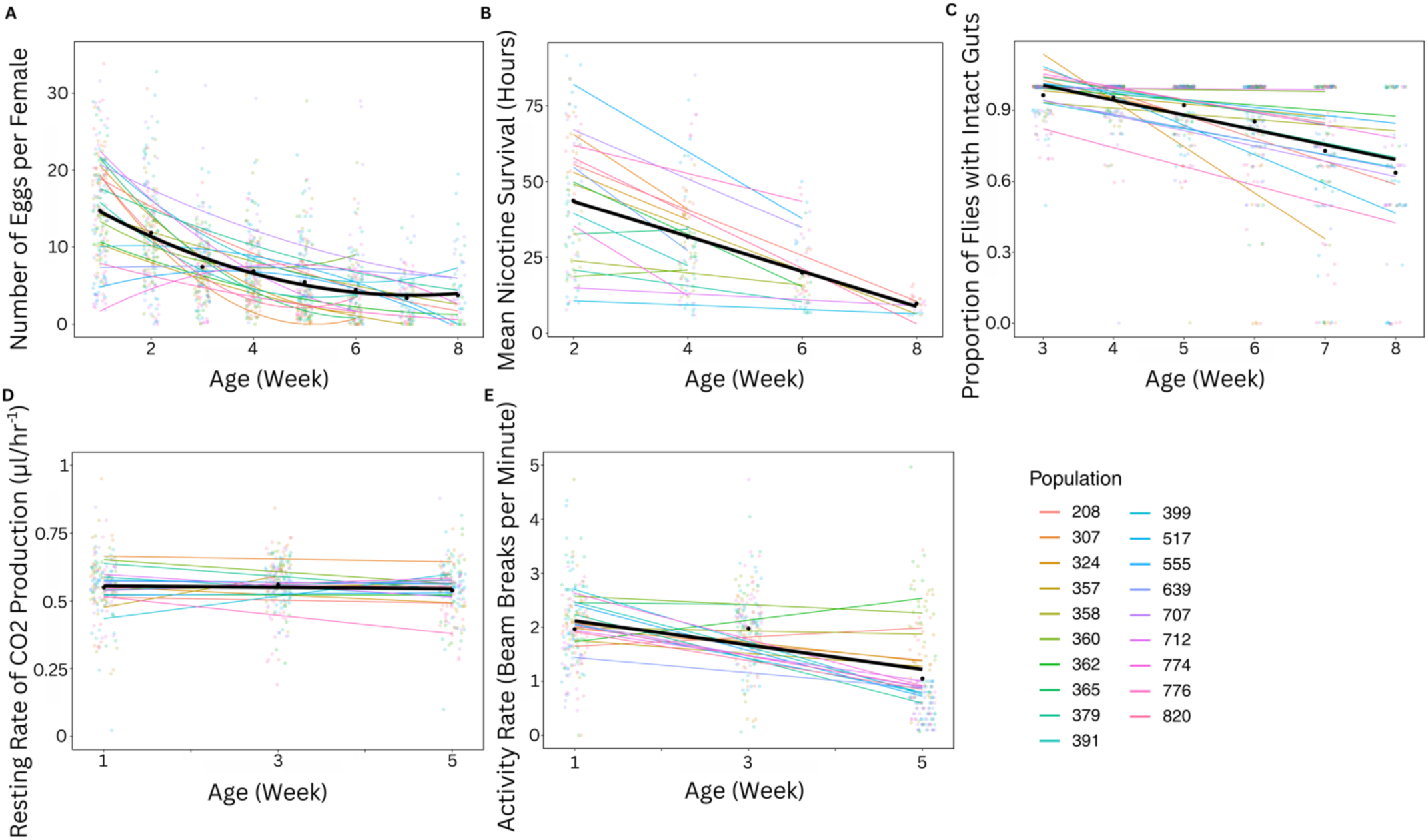
Changes in physiological parameters with age varies significantly between genotypes. Phenotypic assays were performed repeatedly across the lifespans of flies from 19 DGRP lines to measure their decline with ageing. Each phenotype was chosen as a proxy for a different physiological system: (A) egg counts were performed to measure reproductive output; (B) nicotine survival assays were used to measure fat body/ oenocyte detoxification function; (C) smurf assays were performed to measure gut integrity; (D) rate of carbon dioxide production (adjusted for activity and mass), *V̇_C_*_02_, was used as a measure of metabolic rate, and; (E) activity assays were used to measure neuromuscular function. The genotype average for all phenotypes declined with age (Anova Type III: p < 0.01, Supplementary tables S2-S5), except for CO_2_ production (Anova Type III: p = 0.85, Supplementary Table S6), which remained stable with age. Black dots and black lines represent the average effect across genotypes, while coloured dots and lines represent data for individual genotypes. Genotype significantly modified the decline of each physiological parameter (Anova Type III: p = < 0.01, Supplementary tables S2-S6).

Patterns of decline in all assays were significantly different across genotypes (p = < 0.01). Egg production, gut integrity and nicotine survival either declined or remained stable with age. Activity rate and CO_2_ production decreased in some lines but remained stable or increased in others.

### Changes in physiological parameters occur heterogeneously within, and between, genotypes as they age

We found that patterns of decline across physiological parameters differed between DGRP lines (Figure 3). Within each genotype, phenotypes either increased, decreased, or remained stable with age. Furthermore, the identity of those parameters that decreased or otherwise with age differed between lines. For instance, line RAL324 showed strong signs of gut integrity decline with age while line RAL391 did not. Lifespan is often used as a proxy measure of ageing, however by doing this, details regarding underlying physiological decline can be missed. To assess whether lifespan is representative of the decline in each phenotype with age, mean lifespan for each of the 19 DGRP lines was compared against patterns of decline in each of the 5 physiological parameters. Declines in both activity rate and reproduction were found to associate with lifespan, while declines in the other phenotypes did not (Anova Type III: p < 0.01, Supplementary Tables S7 - S11).

**Figure 3.**
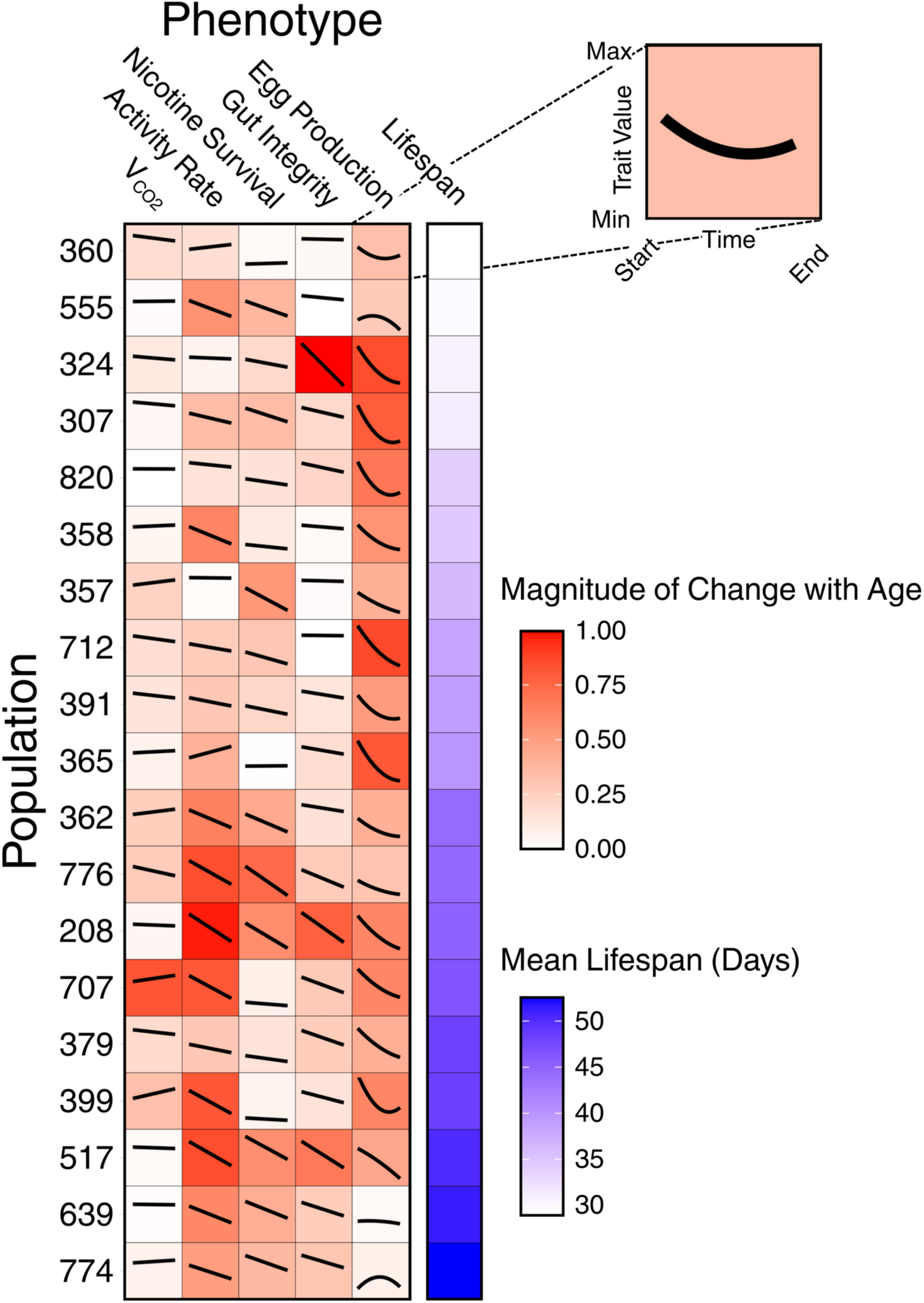
Changes in physiological parameters occur heterogeneously within, and between, genotypes as they age. Red-scale heatmap shows the magnitude of patterns of decline in several physiological metrics, each associated with different organ system functions, for 19 DGRP lines. For each panel, a regression of the normalised phenotypic response (scaled from minimum trait value to maximum trait value across all lines) over time (re-scaled to represent the time from first measurement to last) is shown (see magnified view of top right square). Between genotypes, the magnitude of physiological decline differs significantly with phenotype (Anova Type III: p = <0.01, Supplementary Table S7 - S11). The adjacent blue-scale heatmap represents the mean lifespan of each line.

### Declines in activity rate and reproduction associate with lifespan

While declines in activity did associate with lifespan, there may be more complex patterns of decline which are also able to predict lifespan. To address this, Best Linear Unbiased Prediction (BLUP) slope estimates were obtained for each phenotype change for each genotype to account for uncertainty in individual estimates, and these values were used to perform a Principal Component Analysis (PCA) (Figure 4). The PCA revealed one group of 10 DGRP lines that clustered slightly separately from the other lines, suggesting they have a similar pattern of physiological decline that is distinct from the remaining lines. This cluster had, on average, lower PCA1 scores than the remaining lines, driven primarily by the activity and reproduction phenotypes, indicating they had similar age-related changes in these traits. The remainin DGRP lines were spread across the PCA plot, showing large variation across PC2.

**Figure 4.**
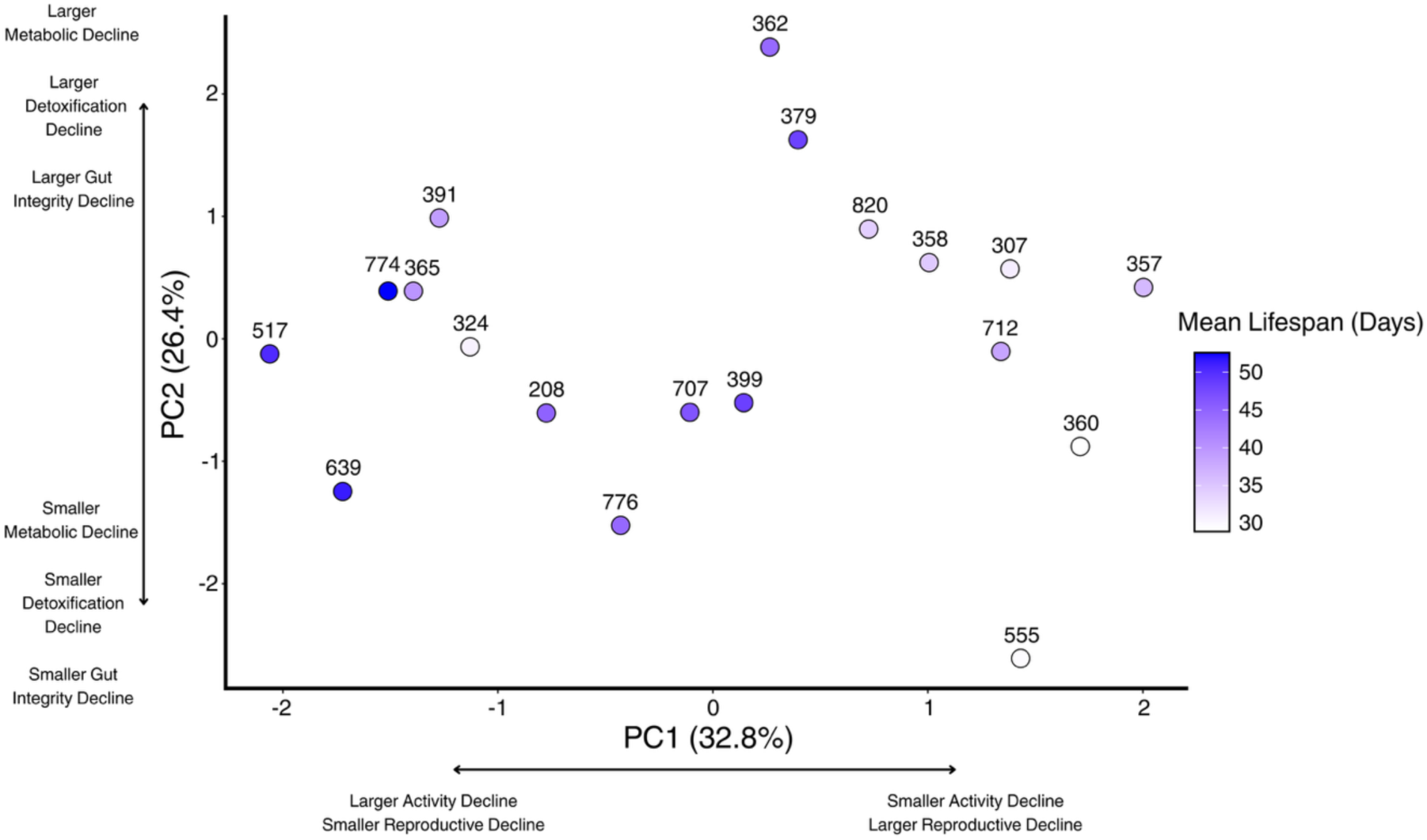
Declines in activity rate and reproduction associate with lifespan. Principal component analysis (PCA) of the rate of decline in egg production, gut integrity, nicotine resistance, *V̇_C_*_02_, and activity rate, reveals no clustering of samples that can explain lifespan variation between lines (point colour scale). PC1 and PC2 combined explain 59.2% of variance in the data. Number above each point represents the DGRP genotype label.

When lifespan outcomes were overlaid onto the points in the PCA, there was an association between time to death and PC1 (p < 0.01, supplementary table 12), suggesting that lifespan can be predicted by rates of activity and reproductive decline.

### Rank order of decline severity of each phenotype differs significantly across DGRP lines

Rather than clustering by variation, we next looked for patterns of physiological decline that might associate with length of life. For each phenotype, we standardised its measured outcomes to a mean of 0 and standard deviation of 1, after which its slope of decline in each genotype was calculated. Using this slope, a rank order of decline severity for each genotype was produced (Figure 5), wherein a more negative slope was considered to represent more severe decline. The rank order of decline severity varied substantially among phenotypes, with Kendall’s coefficient of concordance indicating no agreement across DGRP lines (p = 0.99, supplementary table S13). A group of 3 lines (RAL357, RAL307, and RAL391) and a group of 2 lines (RAL639 and RAL774) shared an identical rank order of decline severity, but these had significantly different lifespan outcomes, ranging from 31 days to 39 days on average. Across lines, each phenotype was represented in rank 1 for at least one line. These data show that the order in which each physiological parameter declines with age is different across lines and is not predictive of final lifespan outcomes.

**Figure 5.**
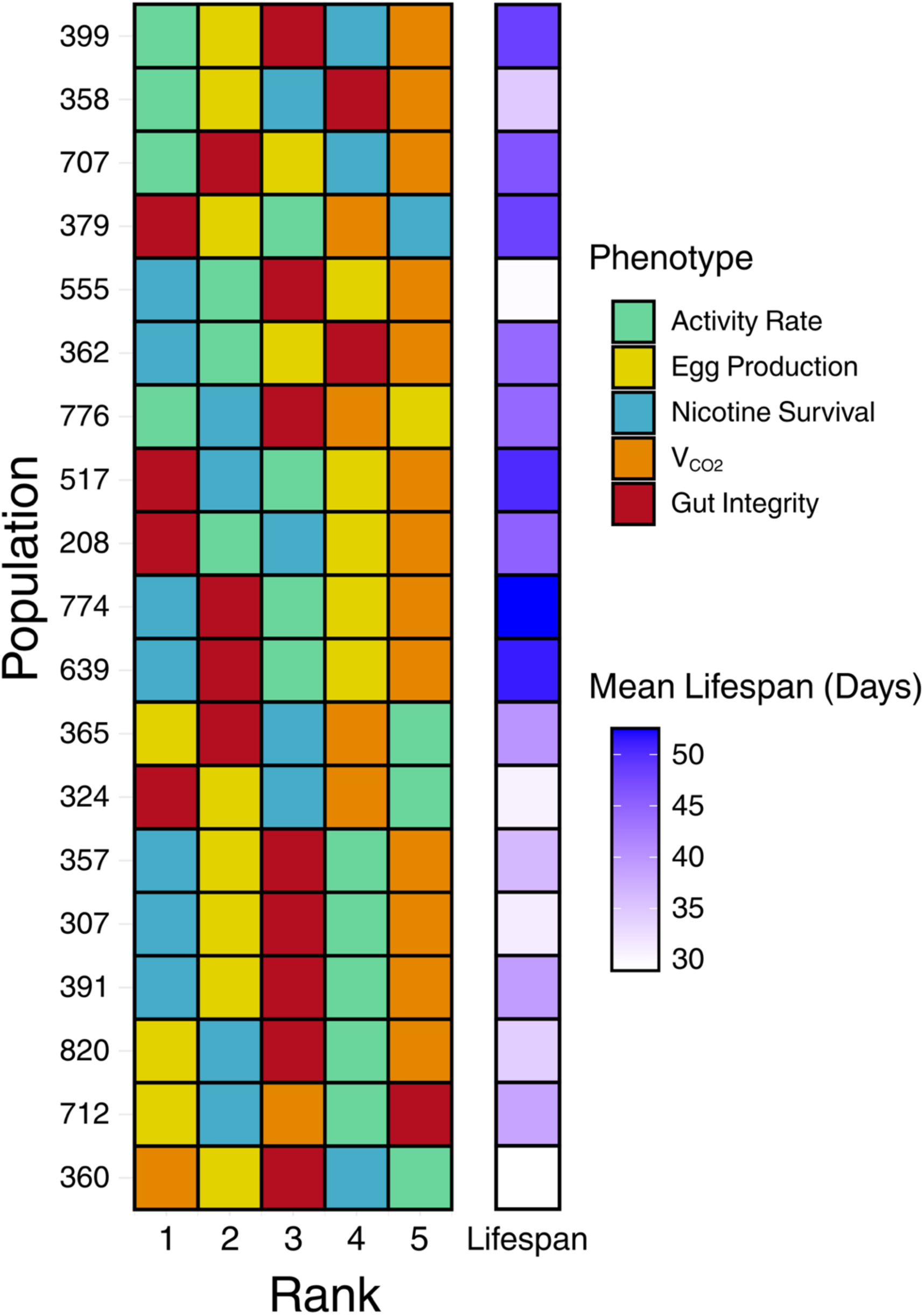
Rank order of decline severity of each phenotype differs significantly across DGRP lines. After standardising the data for each phenotype so that it had a mean of 0 and standard deviation of 1, the slope of decline in each genotype was used to rank the order of decline severity in each of its ageing-related phenotypes (where rank order 1 = most negative slope, to rank 5 = least negative slope). Panel colour represents phenotype. Data was centred so that it had a mean of 0 and standard deviation of 1. Population order in the figure was determined based on pairwise distances among populations using Euclidian distance in R. Rank order was significantly different across DGRP lines (Kendall’s W = 0.14; p = 0.99, supplementary table S13).

## Discussion

It is known that traits associated with ageing, such as lifespan, differ between genotypes (23, 32). While lifespan is a common proxy measure of ageing in lab model organisms, largely due to its relative ease of measurement, underlying this metric is a sequence of physiological decline that leads to the point of death (24, 33). Implicit in the use of lifespan as a proxy measure of ageing is the assumption that individuals with similar lifespans also age in a similar way (24). We found that, on average across genotypes, reproduction, detoxification, activity rate, and gut integrity all declined with age, whereas resting metabolic rate did not significantly decline with age. We found associations between declines in reproduction and activity rate with lifespan. However, we also observed substantial variation in patterns of phenotypic decline across lines, even in individuals with the same or similar lifespans. This means that genetic background should be considered when designing interventions for ageing.

Our data showed that declines in activity and reproduction had a significant association with lifespan, suggesting that individuals with lower activity decline and larger reproductive decline live longer, while those with larger activity decline and lower reproductive decline live shorter. The association between reproduction and lifespan is one that is well established (short lived individuals maximise reproduction early in life compared to longer lived individuals) and thus was expected (34). However, it is unlikely that the decline in reproduction itself is driving shorter lifespans, rather, resource allocation or antagonist pleiotropy mediate both phenotypes simultaneously (35, 36).

While the association between activity and lifespan was also expected, our data, combined with previous work, suggest that it is an imperfect one. It has previously been suggested that although the two traits are associated, lifespan and activity are regulated by different mechanisms and so can be uncoupled (37). This uncoupling is further evidenced by the fact that the relationship observed between these two traits on average across lines was not consistent – some longer lived DGRP lines had a relatively small decrease in activity with age, while some short-lived lines had relatively large age-related activity decline.

Reflecting this imperfect relationship, PC1, driven largely by declines in activity and reproduction, explained a relatively small (32.8%) amount of variation present in our PCA. This suggests that these traits alone do not wholly explain the patterns of age-related physiological decline observed in our data and that other traits, such as gut permeability, metabolic rate, and detoxification also play a significant role. Taken together, although significant interactions between reproductive decline, activity decline, and lifespan were observed, it is unlikely that these traits alone are sufficient to predict lifespan. This further strengthens the idea that individuals in a population do not always follow population trends and highlights the variation in phenotypic decline present in the DGRP.

Previous research has established that metabolic rate does not always associate with lifespan, and this was supported by our data (38, 39). However, traits like improved detoxification ability is common in longer lived individuals, yet we did not find any correlation between this trait and lifespan (40). Previous studies have also shown that the relationship between lifespan and detoxification can become uncoupled. Reduced insulin signalling is associated with both increased lifespan and increased detoxification, however, null mutants of a key mediator of xenobiotic resistance under reduced insulin signalling can affect detox while leaving lifespan untouched (41). This points to a model in which variation in the IIS pathway acts through distinct downstream mediators to influence detoxification and lifespan. Differences in how these mediators respond to increased insulin signalling may play a role in the variation in phenotypic decline observed across all our measured traits.

We found no correlation between length of life and likelihood of smurfing. Recent studies suggest that some form of measurable gut decline occurs preceding death in *Drosophila*, which may also be true for other species (13, 42, 43). Interestingly, several DGRP lines did not exhibit any gut decline throughout life in our assays, and in several other DGRP lines not all individuals smurfed at the same time. There are several possible reasons for this. First, our periodic sampling could easily have missed individual associations between gut decline and time of death, which is generally thought to be ∼48h in *Drosophila* (44). Secondly, it is possible that gut decline occurs in a way that is not captured by the smurf assay meaning that other critical aspects of gut related decline may still occur and not be detected in our experiments (45, 46). Finally, certain genotypes may not express any physiological decline in the gut before death. However, previous smurf assays have been performed in a relatively small number of genetic backgrounds, so there is a possibility that genotypes exist wherein flies do not smurf which have not been discovered as they are yet to be tested (14, 19).

A key strength of the DGRP is that each line represents a single genotype, and thus by comparing phenotypes between lines it is possible to assess the genetic architecture underlying phenotypic variation (20). This has been met with limited success for lifespan, which despite having substantial heritability has proven difficult to map to specific loci (23). This may be the case for two reasons. First, many DGRP lines are short lived because inbreeding can expose deleterious mutations that shorten lifespan (32). However both here and in other studies, lifespan has been shown to vary substantially among the DGRP lines, which is also a feature of the genotypes represented by different individuals in outbred populations (23, 47). In this way, outbred populations can be approximated by the diversity between lines. Thus, using the DGRP to assess variation in ageing-related phenotypic heterogeneity can represent the inter-individual variation in ageing-related phenotypes in other systems (48).

Secondly, lifespan may be the wrong phenotype to map. Our results suggest that genetic variation may act primarily by modulating susceptibility to age-associated physiological decline, from which lifespan is indirectly determined. Thus, isogenic lines may be more informative for identifying genetic variants that influence vulnerability to specific forms of organ-level functional decline during ageing, rather than the overall ageing process.

We observed that all except 5 DGRP lines exhibited a unique order of the magnitude by which their phenotypes declined. This variation in age-related physiological decline across genotypes may arise from differences in the activity of conserved signalling pathways known to modulate lifespan, such as TOR and insulin signalling. These pathways can act in an organ-specific manner, meaning that variation in their rate of signalling may differ between tissues with age and this may modify tissue-specific decline between genotypes (49, 50).

Currently, methods to increase healthy ageing, including those which work to alter TOR signalling, are largely based around treatments that work across a population on average, meaning that while some individuals may garner a benefit to lifespan from a given treatment, others may receive no benefit or be affected negatively (51). These “population average” treatments neglect inter-individual genetic background differences in the way that signalling changes and phenotypes decline, which in turn limits our understanding of ageing at the individual level (52). If the physiological decline underlying lifespan is different across individuals, then treatments aiming to improve healthy ageing should reflect this.

Given our findings, we propose that genetic background itself should be considered a key parameter for how ageing should be treated. The result is an approach to extending healthy life that is personalised and thus potentially more effective than current methods. Future work should aim to assess how genetic variation between lines affects the tissue specific activities of signalling pathways like TOR and insulin, and if this can be predictive of the tissues to fail first during ageing, to provide a mechanistic basis for personalised interventions that target genotype-specific patterns of tissue decline (53).

## Materials and Methods

### Fly Stocks and husbandry

The 19 DGRP lines used in the study were obtained from the Bloomington *Drosophila* Stock Center (20). Stocks were maintained at 18°C, 60% humidity and a 12:12h light/dark cycle. Before use in experiments, stocks were transferred from 18°C to 25°C for three generations. Experimental flies were reared from egg to adult at a controlled density on a sugar yeast (SY) diet (54) (Table S14). To standardise mating status, newly emerged adult flies were kept in mixed cohorts for 48 hours following eclosion. Flies were then lightly anaesthetised with CO_2_, and female flies were sorted into vials containing the SY diet, in cohorts of 10 flies per vial, and 10 replicate vials per genotype, for use in lifespan, gut integrity, reproduction, metabolic rate, and activity assays. An additional 15-20 replicate vials per line, each containing ten flies, were set aside for use in an assay that measured resistance to nicotine (details below). Flies were transferred to fresh food every Monday, Wednesday, and Friday, unless otherwise specified. Experimental flies were maintained at 25°C, 60% humidity and a 12:12h light/dark cycle.

We performed a pilot study on two separate isogenic lines to determine if conducting multiple assays on the same cohort of flies resulted in adverse lifespan effects when compared to a cohort of flies that assessed for lifespan alone. There were no significant differences between flies subjected to multiple assays and flies subjected only to lifespan assays for either genotype (Figure S1, Table S15).

### Blue media preparation for gut integrity assays

Gut integrity was measured using fly food that was dyed blue. SY medium was prepared as normal until the addition of sugar and yeast. After these ingredients were boiled, erioglaucine disodium salt (Sigma-Aldrich, 861146) was added to the media at a concentration of 25g/L, mixed thoroughly, and brought back to the boil. Media preparation was then completed as per standard protocol (54).

### Nicotine-laced media preparation for nicotine resistance assay

Free base nicotine (Sigma-Aldrich, N3876) was diluted in absolute ethanol to a concentration of 175mg/ml. Nicotine-laced medium was prepared by aliquoting 100μL of diluted free base nicotine into a vial containing 3mL of cooled SY medium (final concentration of 5.81mg/mL in vials) (55). Laced vials were then kept in a fabric cover at room temperature for 48h to allow the toxin to diffuse throughout the food. Nicotine-laced food was prepared only in sufficient volumes to match what was needed for immediate use (for each 5-day assay) as nicotine loses potency over time.

### Lifespan

Flies were placed into vials containing 3ml of SY medium at a density of ten flies per vial, with ten replicate vials per genotype. Flies were transferred to fresh vials every two to three days, and deaths and censors were recorded in the software Dlife until all flies in the population died (8, 33).

### Gut integrity (smurf) assay

The protocol described in Martins, McCracken, Simons, Henriques and Rera (42) was adapted to generate periodical snapshots of gut integrity. On day 14 from emergence, flies were switched to the blue SY medium (as described above). After 48 hours on blue food, flies were switched back to the standard (non-blue) SY medium and checked for “smurfing” (42). This phenotype is characterised by the blue dye escaping the gut lumen and entering the body cavity, subsequently turning the whole fly blue. Flies were considered to have smurfed when the blue dye permeated through the gut into the rest of the abdomen, and starting permeating into the thorax (between a “light smurf” and “smurf” on the ‘smurfness’ scale in Martins, McCracken, Simons, Henriques and Rera (42)) (Figure 6). This process was repeated weekly until all flies were dead.

**Figure 6.**
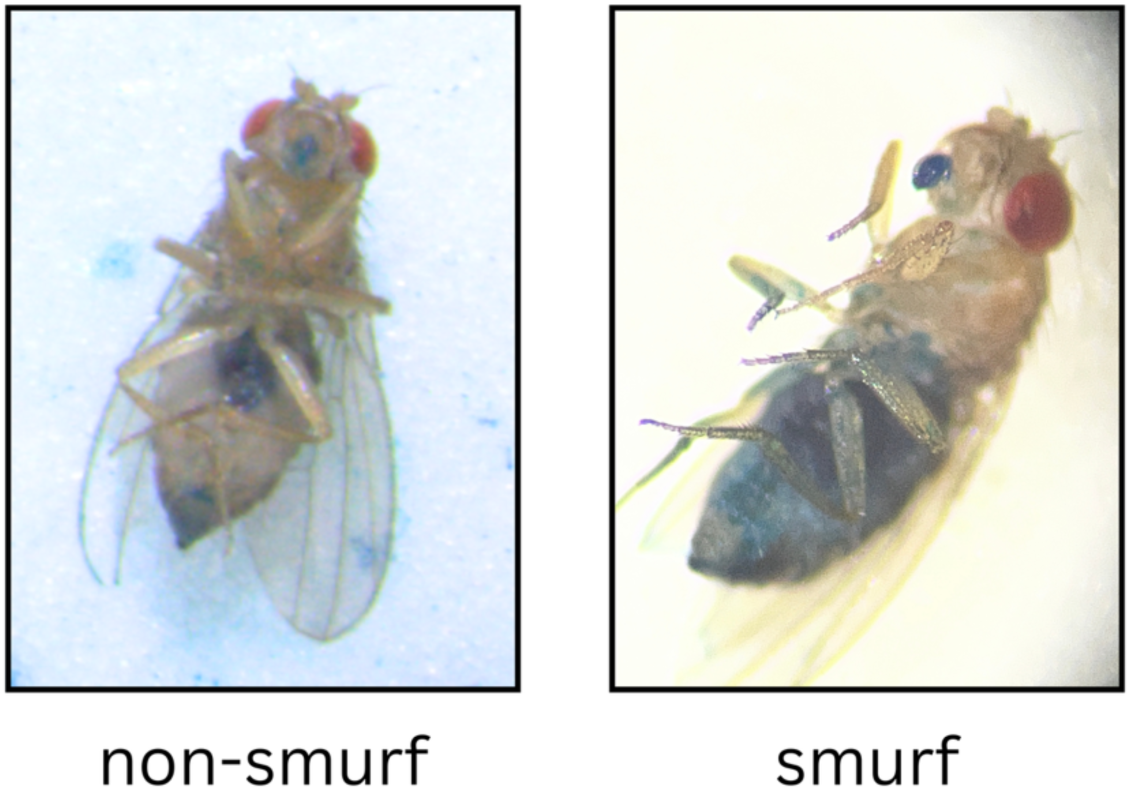
Non-smurf vs smurf. Representative images of flies that were considered “smurfs” and “non-smurfs” in gut integrity assays.

### Nicotine assay

Flies aged on SY food were exposed to nicotine-laced SY food over 5 days, survival was recorded 3 times a day at 7am, 1pm, and 7pm. Any surviving flies following this were censored at the end of the assay period and excluded from analysis. Each 5-day assay used a separate cohort of flies that had been collected and sorted at the same time as the flies used for lifespan and other phenotypic assays. The assay was repeated every 2 weeks for 5 weeks, or until all flies from a population were dead.

### Egg counts

To measure reproductive ageing, flies were maintained on standard SY food and the eggs they laid during a 48h window were counted at intervals throughout life. To count the eggs, flies were transferred to fresh vials and allowed to lay eggs for 48h. Following this period, adults were transferred out, and the vials containing eggs were photographed using a Zeiss Axiocam 212 color camera attached to a Zeiss Stemi 508 dissecting microscope. Fiji (version 1.54p) was used to manually count eggs from the images (56). This was first performed on day 5 and was repeated weekly until all flies were deceased.

### Metabolic rate and activity assay

The rate of CO_2_ production (*V̇_C_*_02_, μl/hr) of adult flies at 25°C was measured as a proxy for metabolic rate using a 16-channel flow-through respirometry system as described in Alton*, et al.* (57) and Alton*, et al.* (30). During the assay, individual flies were confined to separate respirometry chambers that were 65 mm long polycarbonate tubes inserted into Trikinetics *Drosophila* Activity Monitors (DAMs) that allowed for simultaneous measurement of activity (58). Briefly, flies were allowed a 50-minute settling in period at 25°C without food, following which the rates of CO_2_ production and activity were measured for 35 min. The lowest rate of CO_2_ production averaged over a 10 min sliding window was taken as the measure of resting metabolic rate for each fly. Immediately following metabolic rate measurements, the mass of flies was determined. Log_10_-transformed activity data, measured as the average number of times a fly crossed the midpoint of the chamber across the measurement period, (taken from the 10 min CO_2_ production measurement window) and log_10_-transformed mass data were used to adjust metabolic rate data to calculate mass-and activity-independent metabolic rate. (See supplement for detailed methods). Activity data taken from the final 30 min of the measurement period was used to calculate activity rate. Metabolic rate and activity were first measured on day 4 from emergence and subsequently measured once every two weeks for 5 weeks.

### Statistical analysis

All statistical analyses were performed using R version 4.6.1 (R Core 59). Lifespan was modelled using a Cox proportional-hazards implemented with the survival package (60). Significance of effects were evaluated using Type II Anova implemented with the car package (61). Lifespan data was plotted using the survminer package (62).

Egg counts, smurf counts, *V̇_C_*_02_, activity rates, and mean nicotine survival were all analysed individually using linear mixed-effect models implemented with the lme4 package, and the significance of effects were evaluated using Type III Anova’s (63). For each phenotype’s model, block was included as random effect. In addition, log_10_-transformed mass and log_10_-transformed activity were included as continuous covariates in the model for *V̇_C_*_02_. These data were visualised by generating a heatmap using ggplot2, and overlaying trendlines using the patchwork package (64)

Best linear unbiased predictor slope values were found for each phenotype using base R and the lme4 package, and these values were used to generate the PCA. Base R was used to plot the PCA. Associations between the principal components and lifespan were analysed using linear models, and the significance of effects were evaluated using a Type II Anova.

To generate the rank order of decline severity, all measured outcomes were standardised to a mean of 0 and standard deviation of 1, after which slope was generated for each genotype / phenotype combination over time. For each genotype, phenotypes were ranked according to slope, with the most negative slope assigned rank 1 and progressively less negative slopes assigned higher ranks. Kendall’s W, implemented using the

DescTools package, was used to compare the rank order of phenotypes between genotypes (65).

Claude Sonnet 5 was used to assist with data analysis, data visualisation, and manuscript readability.

## Acknowledgments

We would like to thank Professor Craig White for sharing his equipment for measuring metabolic rate and activity. We would also like to thank Louise Cardamone and Lina Vo for their assistance in data collection, and Tahlia Fulton for her assistance with data visualisation. C.K.M was funded by the Australia Research Council (DP250101863). L.A.A. was funded by the Australian Research Council (DP220103421 and FT250100816). This research was supported by the Commonwealth through an Australian Government Research Training Program Scholarship (DOI: https://doi.org/10.82133/C42F-K220).

## Notes

https://doi.org/10.26180/33582502

### Competing Interest Statement

The authors have declared no competing interest.

