## Supplementary Information for "Genetic Variation in Organ System Decline Across the *Drosophila* Genetic Reference Panel"

### Supporting Information Text

#### Methods for Metabolic Rate Measurements

The metabolic rates of individual flies at 25°C was measured by indirect calorimetry. We measured rates of $CO_2$ production ($\dot{V}_{CO_2}$, μL $h^{-1}$) of individual flies using a positive-pressure, flow-through respirometry system that had 16 independent channels that allowed for continuous measurement of $CO_2$ concentration through each channel. To account for variation in metabolic rates associated with activity and body mass, we measured fly activity while inside respirometry chambers using *Drosophila* activity monitors and weighed flies immediately after measuring metabolic rate and activity. When seven lines were phenotyped in a day, we measured 60-64 flies (9-10 flies per line) a random order across four measurement blocks (16 flies per block). For each line, we measured one fly from each vial. Flies that had deformed wings were not chosen for measurement.

To prepare flies for respirometry, a fly selected haphazardly was aspirated from its vial using a mouth-operated pooter with an in-line HEPA filter (Whatman$^{TM}$ HEPA-VENT Disposable Filter) and transferred to one of 16 unsealed respirometry chambers. Each chamber was a polycarbonate plastic tube (5 mm outer diameter x 65 mm length, PPT5x65 Monitor Tube, Trikinetics, Waltham, MA, USA) that had a 10-mm length of foam at each end to contain the fly. This step was repeated until each chamber contained a fly. Each chamber was then inserted into one channel of two *Drosophila* activity monitors (DAM2, Trikinetics, Waltham, MA, USA) with eight chambers in each DAM. Each DAM was then placed inside one of two temperature-controlled cabinets (Refrigerator Incubator, Model MLi125, Nuline Refrigeration, Springvale, Victoria, Australia) where flies were left in the dark at 25 ± 1°C without access to food for ~50 min.

After the 50-min settling time, the chambers were connected to the respirometry system. Dry $CO_2$-free air was pushed through the respirometry system using a purge gas generator (PG14L, Peak Scientific Instruments Ltd., Inchinnan, Scotland, UK) that removed $CO_2$ and water vapour using pressure swing adsorption. The flow rate of air through each of the 16 independent channels of the system was regulated nominally to 50 mL $min^{-1}$ by one of 16 mass flow controllers (Aalborg, Model GFC17, Orangeburg, NY, USA). The volumetric flow rate produced by each flow controller was measured every three weeks prior to starting measurements for a new generation using a Gilian Gilibrator-2 NIOSH Primary Standard Air Flow Calibrator with a low-flow cell (Sensidyne, LP, St Petersburg, FL, USA) and corrected to Standard Temperature Pressure (STP, 101.3 kPa and 0°C). After the flow controller, the air passed through a humidifying chamber (a 10-ml plastic syringe containing four wet cotton balls) before flowing through a respirometry chamber containing an individual fly. The excurrent air from the respirometry chamber then flowed through one channel of eight two-channel infrared $CO_2$/$H_2O$ gas analysers (LI-COR, Model LI-7000, Lincoln, NE, USA) operating in reference estimation mode. The gas analysers measured $CO_2$ concentrations at a resolution of 0.03 ppm and a frequency of 1 Hz and were calibrated every three weeks prior to starting measurements for a new generation with precision span gases (4.2 and 9.8 ppm $CO_2$, BOC Gas, North Ryde, NSW, Australia).

The fractional $CO_2$ concentration of the excurrent air ($F_{eCO_2}$) from each chamber was recorded continuously for 30 min. Locomotor activity during this 35-min measurement period was recorded continuously by the DAM as the number of times the fly walked past the midpoint of the tube (note that the tube length available for movement was 45 mm). The fractional $CO_2$ concentration of the incurrent air without the presence of the chamber was measured for 2 min before and after each 35-min measurement block as a baseline, and a linear model was fitted to these data to estimate the $CO_2$ concentration of the incurrent air during the 35-min measurement period ($F_{iCO_2}$). $\dot{V}_{CO_2}$ was then calculated using equation 1 where $FR$ is the flow rate corrected to STP (i.e. 101.3 kPa and 0°C) accounting for water vapor dilution:

$$\dot{V}_{CO_2} = FR\left(F_{eCO_2} - F_{iCO_2}\right) \qquad (1)$$

The first 5 min of data from the 35-min measurement period were discarded to allow flies to resettle and to allow atmospheric $CO_2$ to washout following connection of chambers to the respirometry system. From the remaining 30 min of data, the lowest $\dot{V}_{CO_2}$ averaged over 10 min was taken as the measure of metabolic rate for each fly. To account for background $\dot{V}_{CO_2}$ associated with the chambers themselves, 16 empty chambers were measured at the beginning of the day using the same protocol as for measuring flies. The lowest $\dot{V}_{CO_2}$ averaged over 10 min was taken as the measure of background $\dot{V}_{CO_2}$ for an empty chamber and was subtracted from the $\dot{V}_{CO_2}$ calculated for the flies measured in the same channel.

The activity recorded during the same 10-min period selected for the metabolic rate calculation was taken as the measure of activity for the fly. However, note that because there was about a 30-s delay between the activity recording and the $\dot{V}_{CO_2}$ recording, and because the DAM was always turned off before the respirometry chambers were disconnected, the measure of activity was sometimes recorded for less than 10 min. To account for the slight differences in the duration of the activity measurement among recordings, activity was divided by the duration of the recording. Thus, activity equated to the number of times a fly walked past the midpoint of the chamber per minute (beam breaks $min^{-1}$).

Immediately following a measurement block of 16 flies, the wet mass of those flies was recorded to the nearest 0.01 mg (XS105DU or XSR105DU Analytical Balance, Mettler Toledo GmbH, Greifensee, Switzerland). Flies were weighed by tipping them from their respirometry chamber into a 0.6 ml plastic tube containing a small amount of egg-laying medium coated with yeast paste. After weighing, tubes containing flies were left unsealed by plugging with them with foam. Flies remained in these tubes until the end of the day when all flies were measured.

**Figures**

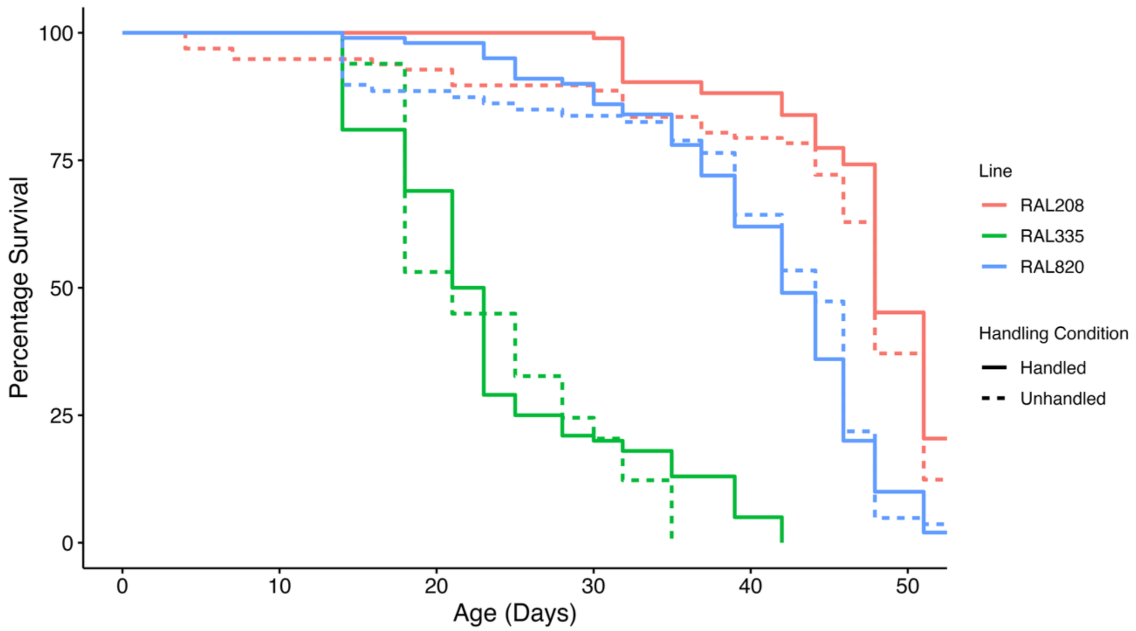


**Figure S1.** Flies subject to phenotyping assays ("Handled") did not have significantly different lifespans to flies that were not ("Unhandled") (Anova Type III: p = 0.277, Table S15)

**Tables**

**Table S1.** Anova (Type II) on Cox Proportional Hazards to evaluate survival differences between genotypes

| | **Chi sq.** | **Df** | **P Value** |
|---|---|---|---|
| **Pop** | 630.75 | 18 | <0.001 |

**Table S2.** Anova (Type III) on linear mixed-effect model to best represent the relationship between egg production, weeks of adulthood (week), and genotype

| | **Chi sq.** | **Df** | **P Value** |
|---|---|---|---|
| **Pop** | 301.00 | 18 | <0.001*** |
| **Poly(week, 2)** | 736.05 | 2 | <0.001*** |
| **Pop:poly(week, 2)** | 380.29 | 36 | <0.001*** |

**Table S3.** Anova (Type III) on linear mixed-effect model to best represent the relationship between gut integrity, weeks of adulthood (week), and genotype

| | Chi sq. | Df | P Value |
|---|---|---|---|
| **Pop** | 35.798 | 18 | 0.007** |
| **Poly(week, 2)** | 81.356 | 1 | <0.001*** |
| **Pop:poly(week, 2)** | 62.128 | 18 | <0.001*** |

**Table S4.** Anova (Type III) on linear mixed-effect model to best represent the relationship between mean nicotine survival time, weeks of adulthood (week), and genotype

| | **Chi sq.** | **Df** | **P value** |
|---|---|---|---|
| **Pop** | 122.327 | 18 | <0.001*** |
| **Week** | 61.941 | 1 | <0.001*** |
| **Pop:Week** | 46.925 | 18 | <0.001*** |

**Table S5.** Anova (Type III) on linear mixed-effect model to best represent the relationship between activity rate, weeks of adulthood, and genotype

| | Chi sq | Df | P value |
|---|---|---|---|
| **Pop** | 24.8662 | 18 | 0.002** |
| **Week** | 2.5282 | 1 | <0.001*** |
| **Pop:Week** | 36.894 | 18 | <0.001*** |

**Table S6.** Anova (Type III) on linear mixed-effect model to best represent the relationship between VCO2, weeks of adulthood (week), mass, activity, and genotype

| | F | Df | P value |
|---|---|---|---|
| **Pop** | 2.8042 | 18 | <0.001*** |
| **Week** | 0.0382 | 1 | 0.85 |
| **Mass** | 9.6521 | 1 | <0.01** |
| **Activity rate** | 8.5927 | 1 | <0.01** |
| **Pop:Week** | 2.1774 | 18 | 0.24179 |

**Table S7.** Anova (Type III) on linear model to best represent the relationship between mean lifespan, gut integrity, and weeks of adulthood

| | **Sum Sq** | **Df** | **F value** | **P value** |
|---|---|---|---|---|
| **Smurf Proportion** | 24.1 | 1 | 0.425 | 0.516 |
| **Week** | 0.2 | 1 | 0.003 | 0.957 |
| **Smurf Proportion:Week** | 5.9 | 1 | 0.104 | 0.748 |

**Table S8.** Anova (Type III) on linear model to best represent the relationship between mean lifespan, activity rate, and weeks of adulthood

| | **Sum Sq** | **Df** | **F value** | **P value** |
|---|---|---|---|---|
| **Activity Rate** | 107.6 | 1 | 1.944 | 0.164 |
| **Week** | 298.8 | 1 | 298.8 | 0.021* |
| **Activity Rate:Week** | 730.7 | 1 | 13.203 | <0.001*** |

**Table S9.** Anova (Type III) on linear model to best represent the relationship between mean lifespan, VCO2, and weeks of adulthood

| | **Sum Sq** | **Df** | **F value** | **P value** |
|---|---|---|---|---|
| **VCO2** | 316.4 | 1 | 5.523 | 0.019* |
| **Week** | 82 | 1 | 1.431 | 0.232 |
| **VCO2:Week** | 78.7 | 1 | 1.373 | 0.242 |

**Table S10.** Anova (Type III) on linear model to best represent the relationship between mean lifespan, egg production, and weeks of adulthood

| | **Sum Sq** | **Df** | **F value** | **P value** |
|---|---|---|---|---|
| **Egg proportion per female** | 41 | 1 | 0.776 | 0.379 |
| **Week** | 1028 | 1 | 19.661 | <0.001*** |
| **Egg proportion per female:Week** | 536 | 1 | 10.255 | 0.001** |

**Table S11.** Anova (Type III) on linear model to best represent the relationship between mean lifespan, VCO2, and weeks of adulthood

| | **Sum Sq** | **Df** | **F value** | **P value** |
|---|---|---|---|---|
| **Activity Rate** | 6.51 | 1 | 0.13 | 0.72 |
| **Week** | 67.03 | 1 | 1.341 | 0.252 |
| **Activity Rate:Week** | 18.64 | 1 | 0.372 | 0.544 |

**Table S12**. Anova (Type II) on linear model to best represent the relationship between mean lifespan and principal components 1-5

| | **Sum Sq** | **Df** | **F value** | **P value** |
|---|---|---|---|---|
| **PC1** | 430.27 | 1 | 11.2484 | 0.005** |
| **PC2** | 10.91 | 1 | 0.2852 | 0.602 |
| **PC3** | 90.97 | 1 | 2.3783 | 0.147 |
| **PC4** | 18.62 | 1 | 0.4867 | 0.498 |
| **PC5** | 37.95 | 1 | 0.9921 | 0.337 |

**Table S13.** Kendall's coefficient of concordance Wt

| **Kendall chi-squared** | **df** | **subjects** | **raters** | **Wt** | **P value** |
|---|---|---|---|---|---|
| **55.046** | 94 | 95 | 4 | 0.146 | 0.999 |

**Table S14.** SY Diet Recipe

| Ingredient | Stock | Quantity per Litre | Supplier | Order # |
|---|---|---|---|---|
| **Water (1st addition)** | | 700mL | Distilled Water | |
| **Grade J3 Agar** | | 10g | Gelita Australia | LR11777 |
| **Bundaberg Sugar** | | 50g | Independent Office Supplies | 620012 |
| **Autolysed Brewer's Yeast Powder** | | 100g | MP Biomedicals | 903312 |
| **Water (2nd addition)** | | 118mL | Distilled Water | |
| **Nipagin** | 100 g/l methyl 4-hydroxybenzoate in 95% EtOH | 30mL | Sigma Aldrich | W271004 |
| **Propionic acid** | | 3mL | Merck | S8309805 |

**Table S15.** Anova (Type III) on Cox Proportional Hazards to evaluate survival differences between flies which were subject to phenotyping assays ("Handled") and flies that were not ("Unhandled")

| | **Chi sq** | **Df** | **P Value** |
|---|---|---|---|
| **Line** | 326.15 | 2 | <0.001*** |
| **Handling** | 1.18 | 1 | 0.277 |
| **Line:Handling** | 1.17 | 2 | 0.556 |
